# Activin E is a TGFβ ligand that signals specifically through activin receptor-like kinase 7

**DOI:** 10.1101/2023.09.25.559288

**Authors:** Kylie A Vestal, Chandramohan Kattamuri, Muhasin Koyiloth, Luisina Ongaro, James A Howard, Aimee Deaton, Simina Ticau, Aditi Dubey, Daniel J Bernard, Thomas B Thompson

**Affiliations:** Department of Molecular and Cellular Biosciences, University of Cincinnati, Cincinnati, OH 45267, USA; Department of Pharmacology and Therapeutics, Centre for Research in Reproduction and Development, McGill University, Montreal, Quebec, Canada; Department of Pharmacology and Systems Physiology, University of Cincinnati, Cincinnati, OH 45267, USA; Alnylam Pharmaceuticals, Cambridge, MA, USA

## Abstract

Activins are one of the three distinct subclasses within the greater Transforming Growth Factor β (TGFβ) superfamily. First discovered for their critical roles in reproductive biology, activins have since been shown to alter cellular differentiation and proliferation. At present, members of the activin subclass include activin A (ActA), ActB, ActC, ActE, and the more distant members myostatin and GDF11. While the biological roles and signaling mechanisms of most activins class members have been well-studied, the signaling potential of ActE has remained largely unknown. Here, we characterized the signaling capacity of homodimeric ActE. Molecular modeling of the ligand:receptor complexes showed that ActC and ActE shared high similarity in both the type I and type II receptor binding epitopes. ActE signaled specifically through ALK7, utilized the canonical activin type II receptors, ActRIIA and ActRIIB, and was resistant to the extracellular antagonists follistatin and WFIKKN. In mature murine adipocytes, ActE invoked a SMAD2/3 response via ALK7, similar to ActC. Collectively, our results establish ActE as an ALK7 ligand, thereby providing a link between genetic and *in vivo* studies of ActE as a regulator of adipose tissue.

**Significance:** Activin E is a homodimeric member of the TGFβ family belonging to the activin subclass. Currently, the signaling capacity of ActE is unknown due to a lack of reliable reagents to study the protein. Here, we demonstrate that ActE acts as a canonical TGFβ ligand that signals through SMAD2/3 in an ALK7-dependent manner, similar to ActC. ActE also utilizes the activin type II receptors, ActRIIA and ActRIIB, to signal and is unable to be antagonized by FS288 and WFIKKN2. This study shows that ActE is a signaling ligand and provides a connection between genetic and *in vivo* studies that links ActE to adiposity.

## Introduction

The Transforming Growth Factor β (TGFβ) superfamily is comprised of over 30 extracellular proteins that play major roles in numerous biological processes, from driving developmental programs in the embryo to tissue regeneration and homeostasis in the adult. Misregulation of TGFβ signaling pathways are associated with a variety of human pathologies including cancer, fibrosis, and pulmonary arterial hypertension. The activins, a subclass of the TGFβ superfamily, have diverse roles in biology, such as regulating reproduction, immune function, inflammation, muscle, and bone mass, along with cell growth and differentiation^1,2^. The activin class is comprised of six primary members that include activin A (ActA), ActB, ActC, ActE; and, the more divergent members, myostatin (GDF8) and GDF11^3^.

The ligands are expressed as precursors that consist of an N-terminal prodomain, important for proper folding and dimerization, and the C-terminal ligand. After synthesis, the prodomain is cleaved from the ligand by furin and two C-terminal ligands are linked via a disulfide bond to complete the dimeric signaling molecule (Fig. 1A)^4^. To signal, ligands bind and assemble a heterotetrameric receptor complex containing two type II and two type I serine/threonine kinase receptors (Fig. 1B). This complex activates intracellular SMAD proteins that function as transcriptional regulators to alter target gene expression. While multiple ligands exist, only 5 type II receptors and 7 type I receptors are available for signaling. The type II receptors are ActRIIA, ActRIIB, BMPR2, TBRII and AMHR2. The type I receptors are collectively referred to as activin receptor-like kinases (ALK) 1-7^5^.

**Figure 1.**
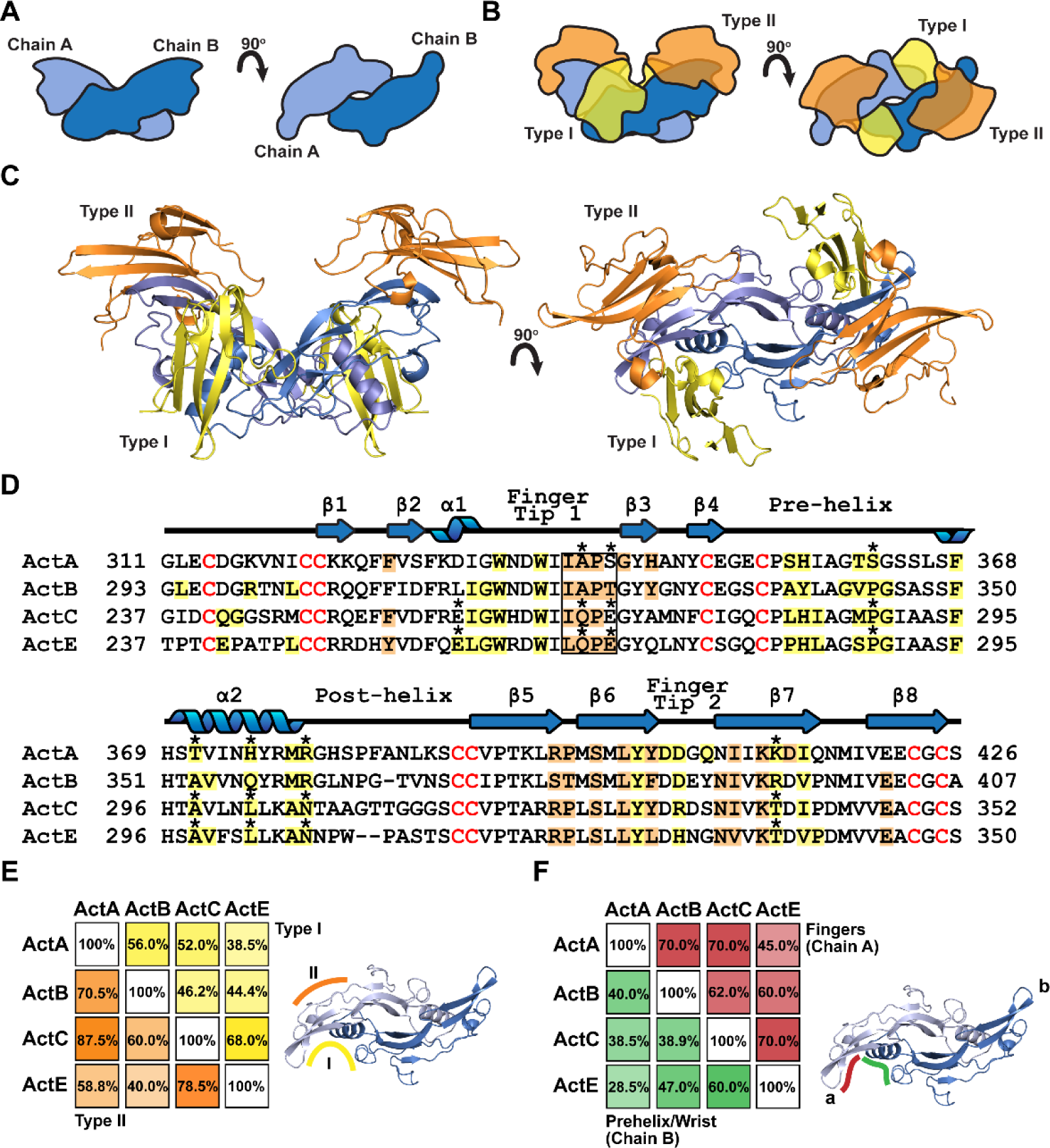
Modeling of the Activin E signaling complex. (A) Schematic representation of a generic dimeric TGFβ dimeric ligand. (B) Schematic diagram of the signaling complex consisting of the dimeric ligand (blue), type I (yellow), and type II (orange) receptors. (C) Overview of the ActE:RIIB:ALK7 ternary complex generated by using AlphaFold 2.3.0. (D) Sequence alignment of the activin subclass annotated based on models of activin ternary complexes (PDB: 7OLY [ActA:RIIB:ALK4], ActB:RIIB:ALK7, ActC:RIIB:ALK7, and ActE:ALK7). Residues are colored if the BSA ≥ 15 Å^2^ – yellow (type I) and orange (type II). Residues that are divergent between the ligands are marked with asterisks. The knuckle region of the ligand central to binding the type II receptor is boxed. (E) and (F) Identity matrix comparing residues across between ligands with (E) comparing the type I and type II receptor and (F) dividing the type I receptor epitope into chain A (fingers, red) and chain B (prehelix/wrist, green)

Ligands of the activin class typically bind with high affinity (nM) to the type II receptors^6^. ActA and ActB are well-studied and primarily bind and signal through ActRIIA and ActRIIB. Multiple crystal structures have demonstrated that the extracellular domain (ECD) of the type II receptors bind independently to each chain of the activin dimer^7,8^. In contrast, the type I receptors exhibit low affinity for activin ligands^9^. Binding of the type I receptor occurs at the dimer interface, creating a composite interface involving contacts with both chains^8,10^. Ligands of the activin class exhibit flexibility at the dimer interface, which partially explains the lower affinity interactions^10^. Type I receptor specificity is variable within the activin class where ActA signals primarily through ALK4, ActB signals through both ALK4 and ALK7 and GDF8 and GDF11 can signal through ALK4 and ALK5^11^. Recently, we established ActC, which was previously thought to be nonsignaling, to signal with high specificity for ALK7^11^.

While the biological roles and signaling mechanisms of multiple members of the activin subclass have been well characterized, ActE is highly understudied. The *Inhbe/INHBE* gene (encoding ActE) was discovered in 1995 from a mouse liver cDNA library and exhibited a liver-specific expression pattern^12–15^. While *Inhbe*^-/-^ knockout mice lacked a significant liver phenotype^16^, subsequent studies showed that ActE overexpression may inhibit proliferation of hepatocytes and pancreatic acinar cells^17–20^. Furthermore, ActE was upregulated in a number of cancers^21–26^.

More recent evidence indicates that ActE has a role in the regulation of adipose tissue. Mouse studies suggest ActE regulates adipogenesis and energy homeostasis^27–30^. In line with this recent human exome-sequencing analyses associated predicted loss-of-function (pLOF) variants in the *INHBE* gene with a more favorable fat distribution and protection from metabolic disorders^31,32^. More recently, genetic studies preformed in mice have shown that overexpression of *Inhbe* increased adipogenesis while *Inhbe* knockout had opposing effects in adipocytes. Interestingly, this study also highlighted that genetic deletion or blocking the type I receptor, ALK7, with a neutralizing antibody inhibited the increase in fat induced by *Inhbe* overexpression, suggesting that the ActE ligand may signal through ALK7^33^.

While these studies present strong evidence, the underlying signaling properties of ActE have not been explored at the protein level. This provokes two important questions: Is ActE a genuine TGFβ signaling ligand, and if so, does ActE directly signal through ALK7? These questions have remained unanswered largely due to the lack of available recombinant ligand. Given the genetic data, we hypothesized that ActE would signal directly through the type I receptor, ALK7. In order to test this hypothesis, we produced recombinant ActE in conditioned media and utilized a variety of cell-based assays to understand ActE signaling mechanisms.

## Results

### Modeling of the Activin E ternary complex

A cross-comparison of activin class ligands shows that it can be divided into three different subgroups: ActA/ActB (64% identity), ActC/ActE (65% identity) and GDF8/GDF11 (90% identity)^12,34^. Given the overall similarity of ActE to ActC, we first wanted to determine how similar the two ligands were at the putative type I and type II receptor binding epitopes. Since there is a lack of structural information on how these ligands bind receptors, we first generated a ligand-receptor model of ActC and ActE bound to a type II and type I receptor. ALK7 was selected as the type I receptor in the model as ActC exclusively signals through this type I receptor. ActRIIB was selected as the type II receptor since it broadly interacts with the known activin ligands, including ActC. To model the ligand:receptor complexes, we utilized AlphaFold Multimer which can build 3D models of protein complexes based on primary amino acid sequences along with information from previously solved structures^35^.

Ternary models of both ActC and ActE bound to the extracellular ligand binding domain of ActRIIB and ALK7 were generated (ActE shown in Fig 1C). The positions of the receptors are consistent with ActRIIB binding to the concave knuckle epitope of each chain and the type I receptor, ALK7, binding at the concave surface of the dimer interface (Fig 1C). The AlphaFold model was essentially identical to a model generated by superimposing individual models of ActE, ActC, ALK7 and ActRIIB onto the ActA ternary complex (PDB: 7OLY)^36^.

Consistent with previous crystallographic studies, the type II receptor buried approximately 750Å^2^ of surface at the knuckle location of ActE while the type I receptor, ALK7, buried 375Å^2^ on chain A and 781Å^2^ on chain B (total 1156Å^2^) (Supp. Tables 1, 2, and 3). Using these models, we wanted to cross-compare the ligand residues at the receptor interfaces. We generated an identity matrix for residues with >15Å^2^ buried surface for the interface with the type I and type II receptors (Fig. 1E). In addition, the type I interface was divided into two components, chain A and chain B, which contain the finger-tip region and prehelix/wrist region, respectively (Fig. 1F).

The analysis revealed that within the different subgroups, sequence identity generally increased when narrowed down to the interfacing residues as compared to the overall sequence identity. For instance, a comparison of the residues at the type II and type I interfaces of ActE with ActC increased identity to 78% and 68%, respectively, suggesting that ActE and ActC may exhibit similar preference for type II and type I receptors (Fig. 1E). Interestingly, ActE has a similar glutamine at the type II interface as observed in ActC (alanine in ActA and ActB, see box in Fig1D). Variations of residues at this location are thought to be important for type II receptor affinity differences ^7,8^. Interestingly, conserved differences between ActE and ActC were also identified in key positions at the type I interface, namely in both the finger-tip and prehelix regions, which both have been shown to be important for type I receptor specificity (see residues marked with an * in Fig1D)^10^. Thus, the modeling demonstrated that ActE and ActC are even more similar at the type I and type II receptor binding sites as compared to their overall identities, consistent with the hypothesis that ActE and ActC may signal through similar receptors, ActRIIA/B and ALK7.

### Activin E induces SMAD2/3 activation through ALK7

Since recombinant ActE is not commercially available, we designed two distinct constructs of full-length ActE, Construct I and Construct II, for mammalian cell expression and to test if ActE could activate TGFβ family receptors (Fig. 2A). Construct I was designed with a Flag-Myc tag on the N-terminus of the prodomain whereas the mature domain was tag-free. Construct II included a Myc and Flag tag on either side of the furin cleavage site which would maintain a Flag-tagged mature ligand and allow for detection by western blot (Fig. 2A). In both cases, the furin site was modified to a potentially more optimal recognition sequence from RARRR to RRRRR. Fig 2B provides a schematic to highlight the expected molecular weights (MW) of various protein species that can arise from incomplete furin processing and the expected differences in species under non-reduced and reduced conditions. The prodomain of ActE contains a single N-linked glycosylation site (* in Fig 2A) which can increase the apparent MW of certain species.

**Figure 2.**
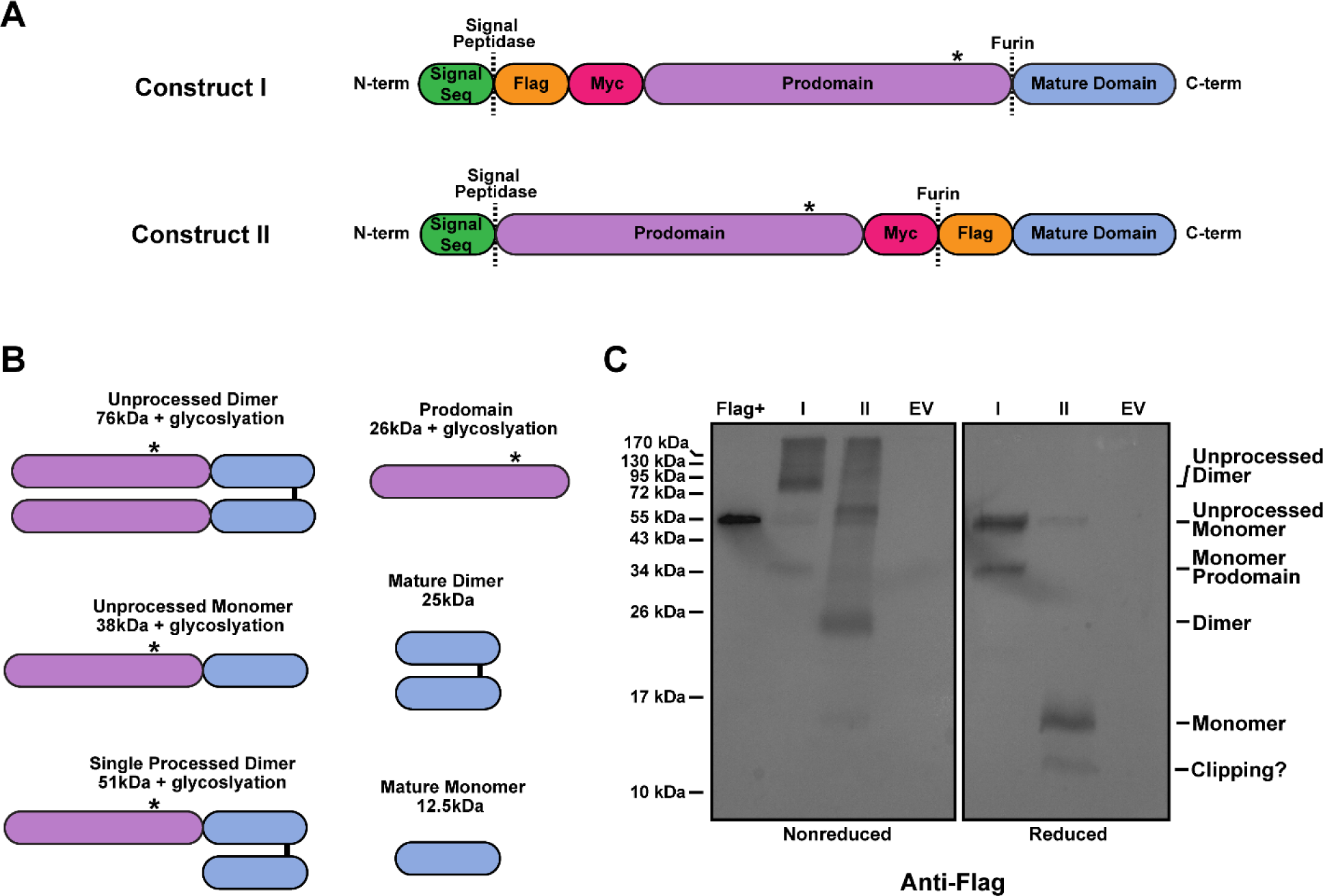
Construct design and expression of Activin E. (A) Schematic of the full-length ActE constructs (Construct I and Construct II) used for mammalian cell expression in ExpiCHO-S cells. Asterisks denote the known N-linked glycosylation site on the prodomain. (B) Schematic of the different processing states of ActE and corresponding sizes (without glycosylation). (C) Western blot analysis (α-Flag) showing expression of Constructs I and II along with empty vector (EV) in ExpiCHO-S media under both non-reducing (left) and reducing (right) conditions. Flag + control is Flag-BAP fusion protein run under non-reducing conditions (left).

Each construct was transiently expressed in Chinese hamster ovary cells (Expi-CHO) and the media was analyzed by an anti-flag western blot (Fig 2C). For Construct I, we observed the majority of ActE as unprocessed dimer (∼80 kDa) with a small amount of prodomain (∼38kDa). Under reducing conditions, we observed an unprocessed monomer (∼55 kDa) along with the prodomain. Due to the lack of a tag on the mature domain, we were unable to detect the mature ligand (25 kDa nonreduced and 12.5 kDa reduced) (Fig. 2C). For Construct II, the major species migrated at 25 kDa and 12 kDa on nonreduced and reduced conditions, consistent with a Flag-tagged ActE dimer (Fig. 2C). For both constructs, furin processing was incomplete resulting in larger MW species representing single and double unprocessed species. Interestingly, there exists a lower MW species that is smaller than the ActE dimer. This may be a clipping product of mature ActE, suggesting that the mature dimer may be sensitive to proteolysis. Expression of Construct II clearly produces ActE dimer while Construct I shows expression of a processed ActE prodomain and is potentially producing mature ActE.

We next determined whether either Construct I or II could produce a signal through TGFβ family receptors. Initially, we used a luciferase reporter assays in HEK-293T cells stably transfected with the (CAGA)12-luciferase plasmid^9,36^. These reporter cells endogenously express only two of the three activin type I receptors, ALK4 and ALK5, but not ALK7, along with both type II receptors, ActRIIA and ActRIIB^9^. We transiently transfected Construct I and II at low (5ng) or high (50ng) concentrations of DNA to test activity, along with control plasmids that express either ActA or ActC^11,37^. As expected, transfected ActA robustly stimulated promoter activity (Fig. 3A). Consistent with previous results, ActC, due to the lack of endogenous ALK7 in HEK-293T cells, did not induce (CAGA)12 promoter activation. Similarly, both ActE constructs showed no (CAGA)12 promoter activity in HEK-293T cells (Fig. 3A).

**Figure 3.**
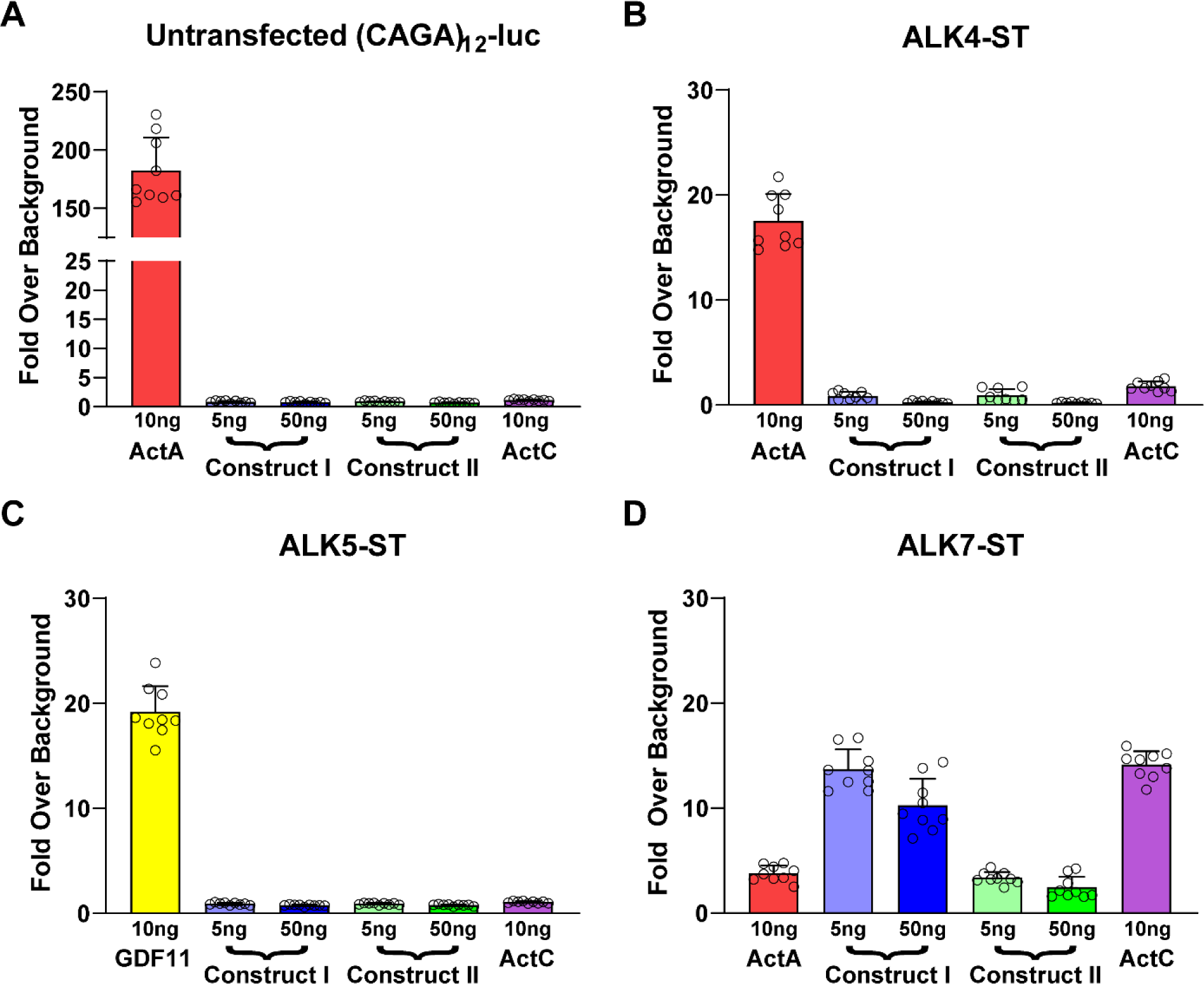
Transient expression of ActE activates signaling through ALK7. (A) CAGA-luciferase reporter assay in HEK293T293 cells in response to transfection of ActA (10ng), ActC (10ng), or ActE Construct I (5ng or 50ng), and ActE Construct II (5ng or 50ng). (B, C, and D) Similar assay to (A) modified by transfection of SB-431542 resistant type I receptors; ALK4-ST (B), ALK5-ST (C), and ALK7-ST (D) type I receptors. All cells were transfected with 100ng total of DNA containing either empty vector (EV) and ligand or EV, ligand, and type I receptor. Data normalized to cells not transfected with ligand DNA constructs (EV alone or EV and type I receptor). Each point represents a technical replicate within triplicate experiments with bars displaying the mean ± SD. In (B, C, and D), cells were treated with 10μM SB-431542 to inhibit signaling activity of endogenous receptors.

To extend this assay to include ALK7 we turned to our previously established system where we transfect in type I receptors with single point mutations (ALK4-ST, ALK5-ST, ALK7-ST) in the kinase domain active site that render it resistant to inhibition by the small molecule kinase inhibitor SB-431542^9,10^. Thus, by treating cells with SB-431542, we can effectively block signaling via endogenous type I receptors and enable unencumbered signaling through the resistant type I receptors. As expected, ActA signaled through ALK4-ST (Fig. 3B) and GDF11 signaled through ALK5-ST (Fig. 3C)^9,10,38^. Similar to ActC, both ActE constructs were unable to stimulate reporter activity in the presence of ALK4-ST or ALK5-ST (Fig. 3B and 3C). Notably, ActE Construct I and to a lesser extent Construct II stimulated the (CAGA)12 promoter in HEK-293T cells transfected with ALK7-ST suggesting that ActE signals through ALK7 (Fig. 3D). Similar to previous results, ActC activated (CAGA)12-luc activity with ALK7-ST while ActA minimally activated the promoter under these conditions^11^.

To further analyze ActE signaling, we tested the signaling potential of conditioned media generated from cells transfected with Construct I or II. We compared ActE conditioned media shown in Fig. 2C to purified protein controls (ActA, GDF11 and ActC) using the same strategy of SB-431542 resistant type I receptors in the (CAGA)12 promoter luciferase assay in HEK-293T cells. As expected, ActA elicited a response in the ALK4-ST luciferase assays whereas ActC did not (Fig 4A). ActE conditioned media minimally induced a response in the ALK4-ST assays similar to the empty vector (EV) conditioned media control (Fig. 4A). Additionally, all conditioned media samples (Construct I, Construct II, and EV) failed to activate the reporter in the ALK5-ST transfected cells similar to ActC and unlike GDF11 which activated the reporter as previously shown (Fig. 4B)^9^. Remarkably, the media conditioned with either ActE Construct I or II activated (CAGA)12-luc activity in the presence of ALK7-ST, similar to ActC (Fig. 4C). EV conditioned media showed no signaling through ALK7-ST (Fig. 4C).

**Figure 4.**
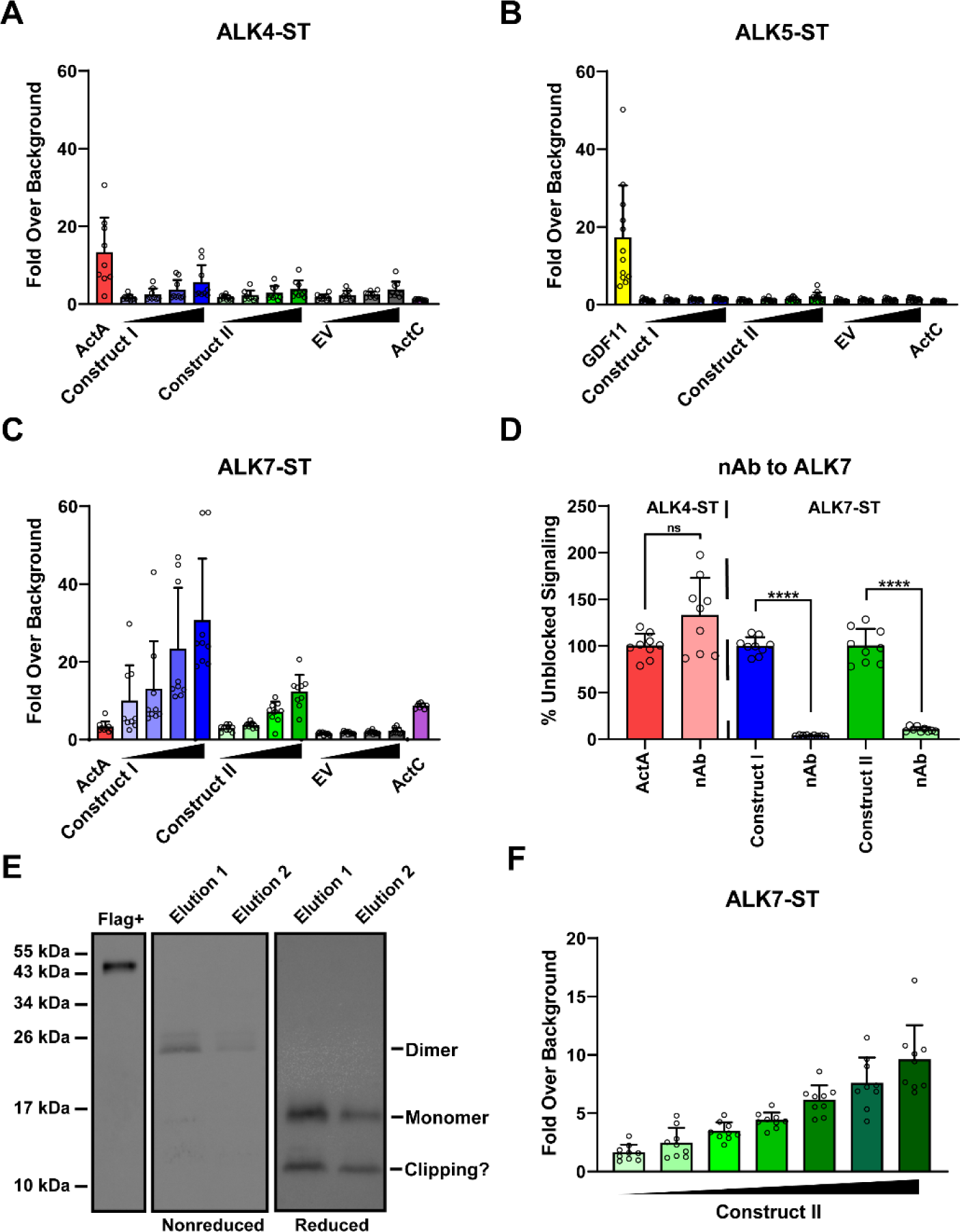
ActE conditioned media signals though ALK7. (A, B, and C) Luciferase reporter assay response to recombinant mature ActA, ActC, or GDF11 (5 nM) along with conditioned media of ActE Construct I, Construct II, or empty vector (EV) in (CAGA_12_)-luciferase HEK293T293 cells transfected with SB-431542 resistant ALK4-ST (A), ALK5-ST (B), or ALK7-ST (C) type I receptors. Media was concentrated 20x and diluted 2-fold for the titration (2.5x, 5x, 10x, 20x). Data normalized to transfected cells with no ligand treatment. (D) Effects of the ALK7 neutralizing antibody (nAb) on recombinant ActA or ActE conditioned media samples on (CAGA_12_)-luciferase promoter activity in cells expressing the designated SB-431542 resistant type I receptor. (ns = not significant) (****P < 0.0001) (****P < 0.0001) (E). Western blot analysis under reducing and non-reducing conditions of elution samples from the purification of flag-tagged mature ActE from construct 2 using flag affinity resin. (F) Luciferase reporter activity of pooled fractions from (E) in (CAGA_12_)-luciferase HEK293T cells transfected with the ALK7-ST. Data normalized to untreated control cells. In (A, B, C, D, and F) each point represents a technical replicate within triplicate experiments with bars displaying the mean ± SD. In (A, B, C, D, and F), cells were treated with 10μM SB-431542 to inhibit signaling activity of endogenous receptors.

These activity data suggest that ActE can signal using ALK7 as its type I receptor. To validate these results, we used an ALK7-specific neutralizing antibody that binds the ECD of the receptor and blocks signaling of ALK7 dependent ligands. Previous studies showed that ActC signaling is abrogated by the anti-ALK7 neutralizing antibody^11^. Addition of the anti-ALK7 antibody robustly abolished signaling of ActE in conditioned media from cells transfected with either Construct I or II (Fig. 4D). The antibody did not affect ActA signaling via ALK4.

Next, we attempted to purify the mature ActE ligand through Flag-affinity purification using conditioned media with Construct II (Flag-ActE). Upon elution, bands were identified at the mature dimer and monomer sizes which were confirmed through western blot analysis directed against the Flag tag (Fig. 4E). The semi-purified Flag-ActE was able to signal through ALK7-ST to activate the (CAGA)12-luc promoter in a dose-dependent manner (Fig. 4F). Collectively, these data show that ActE signals specifically through ALK7, similar to ActC. Since N-terminal tag on ligands can interfere with receptor binding, for subsequent testing we focused on ActE derived from Construct I (tag-free), especially as it exhibited higher activity than ActE from Construct II (Fig. 4C). Unfortunately, we do not yet have the means to purify the untagged ActE and therefore, are reliant on the conditioned media at this time.

### ActE requires activin type II receptors to signal

Along with type I receptors, ligands utilize the type II receptors in order to activate intracellular SMAD signaling. To test whether ActE uses the canonical activin type II receptors, we used type II receptor-Fc constructs (decoy receptors) to block the ligands from interacting with cell surface receptors in luciferase assays (Fig. 5A). We titrated in either ActRIIA-Fc or ActRIIB-Fc (6.25, 12.5, or 25nM) against a constant concentration of ligand (0.62nM) or conditioned media (20x) (Fig. 5B and 5C). ActRIIA-Fc inhibited ActA signaling in a dose-dependent manner while ActRIIB-Fc nearly abrogated signaling at all concentrations in the ALK4-ST assay. In the ALK7-ST assay, ActRIIA-Fc inhibited signaling by ActC in a dose-independent manner while ActRIIB-Fc titration failed to inhibit ActC signaling even at the highest concentration (25nM). Similarly, ActRIIA-Fc was able to inhibit ActE signaling in a dose-dependent manner in the ALK7-ST assay; however, ActRIIB-Fc had no effect on ActE signaling, similar to ActC (Fig. 5B and 5C).

**Figure 5.**
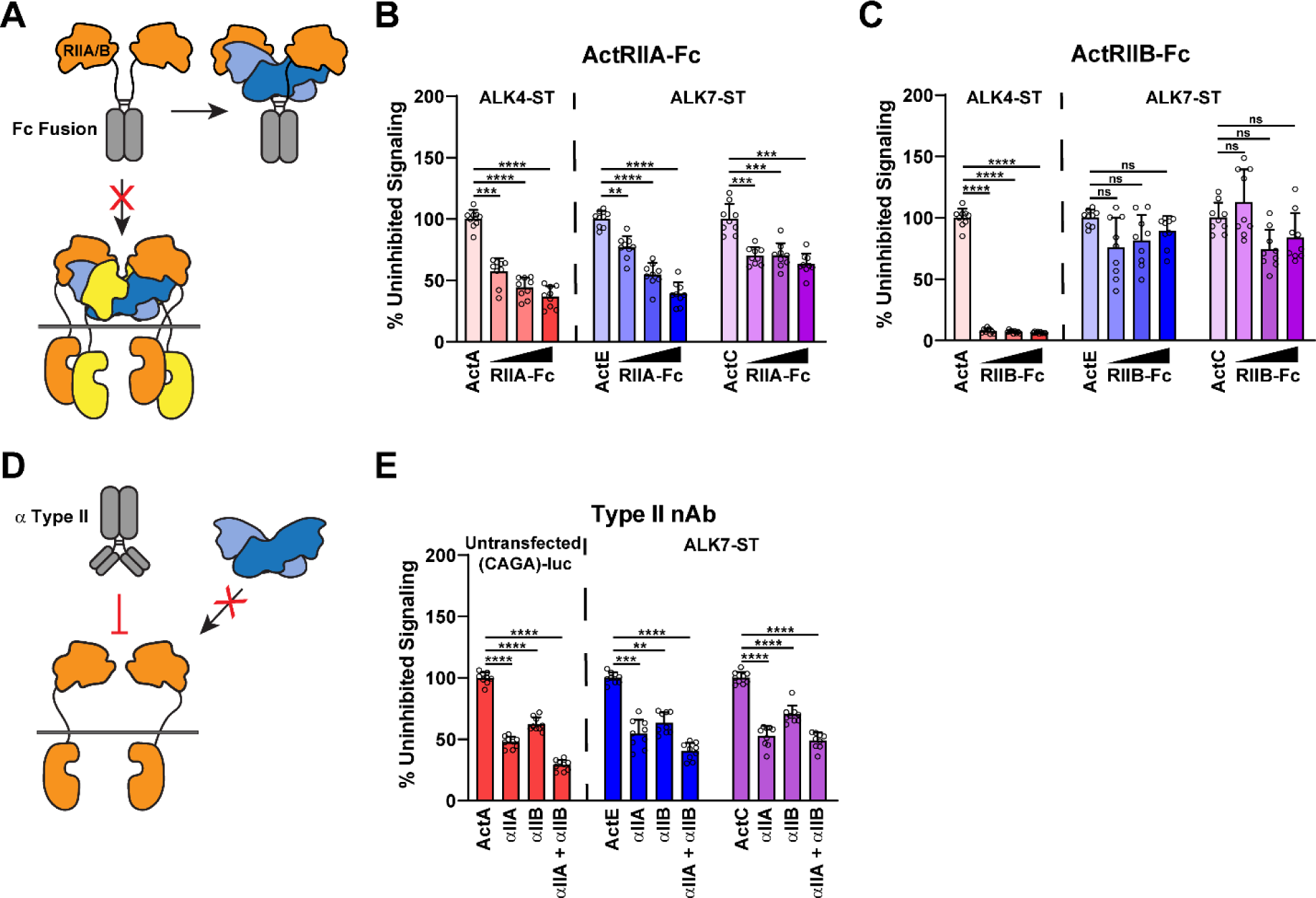
ActE signals via the activin type II receptors. (A) Schematic diagram of activin type II receptor Fc-fusion proteins used as decoy receptors. (B and C) (CAGA_12_)-luciferase HEK293T293 cells transfected with ALK4-ST or ALK7-ST receptor constructs and treated with ActA (0.62nM), ActE conditioned media (20x), ActC (0.62nM), and _10μM_ SB-431542 in the presence of increasing amounts (6.25, 12.5, 25 nM) of either ActRIIA-Fc (****P < 0.0001) (****P < 0.0001) (****P = 0.0002) (B) or ActRIIB-Fc (****P < 0.0001) (P = 0.3132) (P = 0.0549) (ns = not significant) (C). (D) Schematic representation of neutralizing antibodies targeting the extracellular domains (ECDs) of the activin type II receptors. (E) (CAGA_12_)-luciferase HEK293T293 cells either untransfected or transfected with the ALK7-ST type I receptor and treated with ActA (0.62nM), ActE conditioned media (20x), ActC (0.62nM), and _10μM_ SB-431542 in the presence or absence of neutralizing antibodies targeting ActRIIA, ActRIIB, or both (2.5µg/mL) (****P < 0.0001) (****P < 0.0001) (****P < 0.0001) (ns = not significant) . In (B, C, and E) each point represents a technical replicate within triplicate experiments with bars displaying the mean ± SD. Data are represented as percentages of uninhibited signal (B, C, and E).

Next, we examined whether ActE signaling through ALK7 was dependent on the type II receptors. To test this, we added neutralizing antibodies that target the extracellular domains (ECDs) of ActRIIA, ActRIIB, or both receptors (2.5µg/mL) (Fig. 5D). As expected, neutralization of either ActRIIA or ActRIIB significantly decreased signaling by ActA in an untransfected assay with only endogenous receptors. The antibody directed against both type II receptors blocked signaling more potently compared to the single target antibodies. In the presence of ALK7-ST, ActE and ActC signaling was significantly reduced when ActRIIA was neutralized and to a lesser extent when ActRIIB was blocked (Fig. 5E). Altogether, these data suggest that ActE requires the activin type II receptors, with a possible preference to ActRIIA, to signal through ALK7.

### ActE is resistant to antagonism by Follistatin and WFIKKN

Regulation of signaling by extracellular antagonists in a common feature for most TGFβ family ligands. Most ligands of the activin class are neutralized by extracellular antagonists of the follistatin family which sterically block the binding of both the type I and type II receptors^9,37,39–41^. Since we previously established that ActC is not antagonized by FS288, a splice variant within the follistatin family, we hypothesized that ActE might be similarly resistant. As predicted, FS288 had no effect on ALK7-dependent signaling by ActE from conditioned media (Fig. 6A). ActE conditioned media along with EV conditioned media slightly stimulated promoter activity in the presence of ALK4-ST (Fig 4A and 6A). This activity was neutralized by the addition of FS288 (Fig. 6A), suggesting the presence of trace amounts of ActA or ActB in the media.

**Figure 6.**
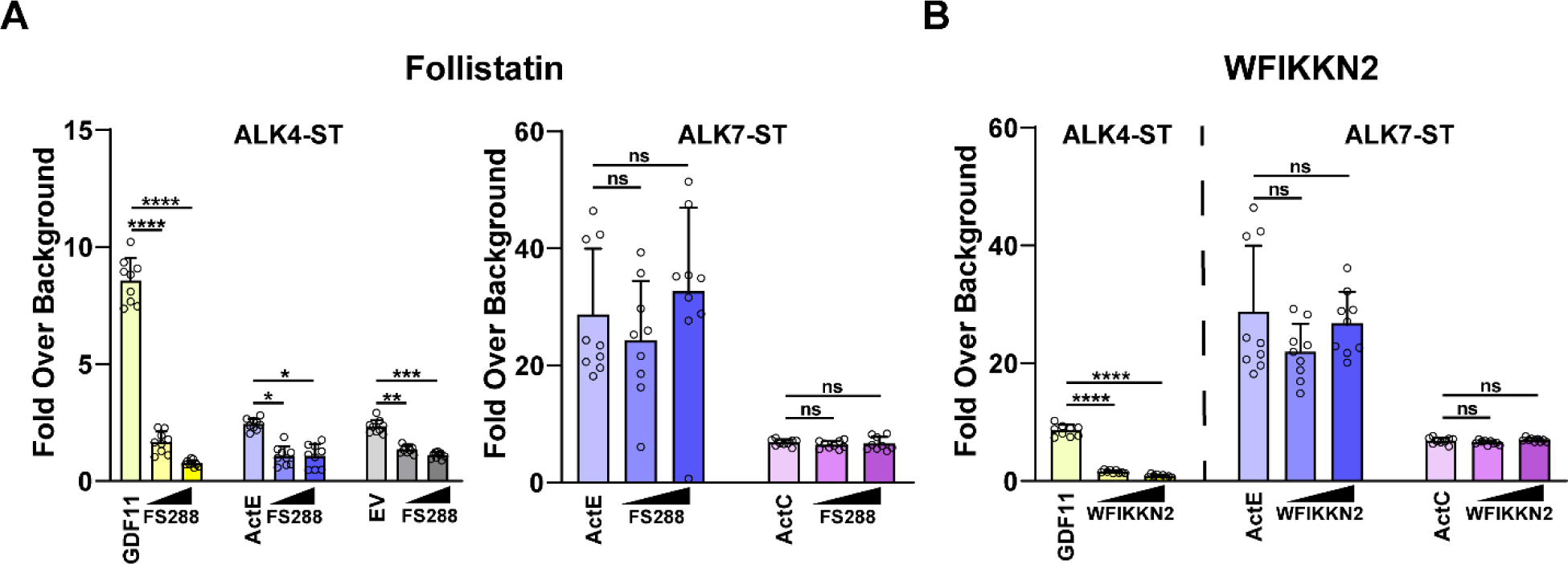
Activin E is resistant to antagonism by Follistatin and WFIKKN. (A) (CAGA_12_)-luciferase HEK293T cells transfected with ALK4-ST or ALK7-ST receptor constructs and treated with recombinant GDF11 (0.62nM), ActE conditioned media (20x), EV conditioned media (20x), and ActC (0.62nM) in the presence or absence of increasing amounts of FS288 (12.5 and 25nM) (****P < 0.0001) (*P = 0.0105) (***P = 0.0004) (P = 0.2746) (P = 0.8552) (ns = not significant). (B) (CAGA_12_)-luciferase HEK293T cells transfected with ALK4-ST or ALK7-ST receptor constructs and treated with recombinant GDF11 (0.62nM), ActE conditioned media (20x), EV conditioned media (20x), and ActC (0.62nM) in the presence or absence of increasing amounts of WFIKKN2 (12.5 and 25nM) (****P < 0.0001) (P = 0.3687) (P = 0.1983) (ns = not significant). Each point represents a technical replicate within triplicate experiments with bars displaying the mean ± SD. Data are normalized to untreated control cells. In (A and B), cells were treated with 10μM SB-431542 to inhibit signaling activity of endogenous receptors.

While follistatin binds and neutralizes several activin class ligands, the antagonists, WFIKKN1 and WFIKKN2, are highly specific for GDF8 and GDF11. Therefore, we tested if WFIKKN2 could antagonize ActC or ActE. As previously shown, GDF11 signaling was nearly abolished at both concentrations of WFIKKN2^42,43^. However, addition of WFIKKN2 did not inhibit ActE or ActC signaling through ALK7 (Fig. 6B). Overall, these data indicate that ActC and ActE are similarly resistant to FS288 and WFIKKN2, which are common extracellular antagonists of the activin class.

### ActE activates SMAD phosphorylation in mature adipocytes

Finally, we wanted to establish that ActE can signal through ALK7 in cells that endogenously express the type I receptor. We, therefore, utilized mature adipocytes from the stromal vascular fraction (SVF) of mouse adipose tissue. We previously reported that these cells express endogenous ALK7 and respond to ActC^11,44^. In mature adipocytes, ActC (recombinant) and ActE (conditioned media) stimulated SMAD2 phosphorylation (Fig. 7A, lanes 3 and 5). These effects were blocked by addition of the ALK7 neutralizing antibody (Fig. 7A, lanes 4 and 6). As in the reporter assays, FS288 did not block ActC or ActE mediated SMAD2 phosphorylation (Fig. 7B, compare lanes 3 vs. 4, and 5 vs. 6). Interestingly, while less potent, the EV conditioned media induced a small amount of pSMAD2 as well (Fig. 7A, lane 1). This was unaffected by the ALK7 neutralizing antibody (Fig. 7A, lane 2) but was reduced by FS288 (Fig. 7B, lane 1 vs. 2). Taken together, the ‘basal’ pSMAD2 in the EV likely reflects the actions of ActA signaling through ALK4 and is consistent with the luciferase reporter assays (Fig. 6A). Collectively, these results show that ActE, like ActC, can activate an ALK7-specific signal that is resistant to the extracellular antagonist, FS288, in adipocytes.

**Figure 7.**
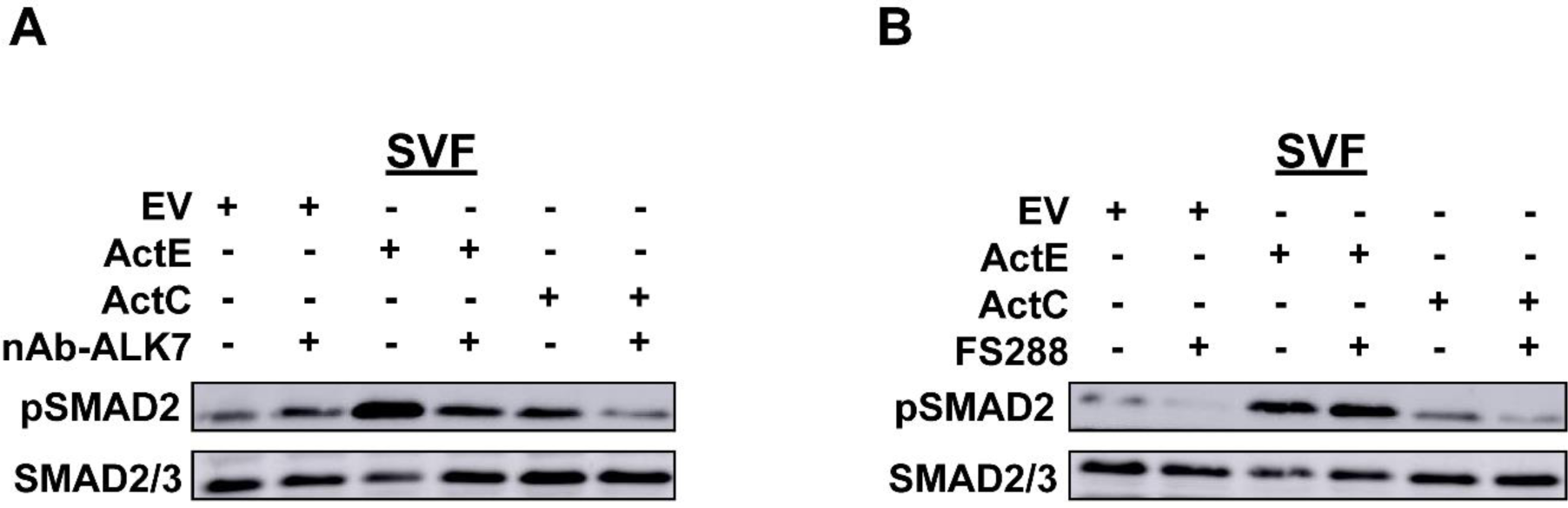
ActE activates phosphorylation of SMAD2 via ALK7 in mature adipocytes. Western blot analysis showing phosphorylated SMAD2 (pSMAD2) and total SMAD2/3 in SVF-derived adipocytes following treatment with EV conditioned media, ActE conditioned media (20x), or ActC in the presence or absence of an ALK7 neutralizing antibody for 1 h. (B) Western blot analysis showing phosphorylated SMAD2 (pSMAD2) and total SMAD2/3 in SVF-derived adipocytes following treatment with EV conditioned media, ActE conditioned media (20x), or ActC in the presence or absence of FS288 for 1 h. Data shown are representative of two independent experiments.

## Discussion

The signaling mechanisms of most activin class members, ActA, ActB, GDF8, and GDF11, have been well characterized^9–11,37,39,42^. Each member possesses differences in specificity for both the type II (ActRIIA and ActRIIB) and type I receptors (ALK4, ALK5, and ALK7). For example, ActA has little preference for the type II receptors while primarily only using ALK4^10,36^. In contrast, GDF11 has a preference towards ActRIIB while being promiscuous in its type I utilization, signaling via ALK4 and ALK5 and modestly through ALK7^9^. Recently, we discovered that ActC is a bona fide member of this class as it stimulates the SMAD2/3 branch of the TGFβ pathway through activation of ALK7^11^. However, ActC is distinct in that it has a low affinity for type II receptors and is surprisingly resistant to follistatin antagonism.

Prior to this study little information on the potential of ActE to function as a signaling molecule has been shown. Past studies have speculated that ActE may utilize ALK7 as its type I receptor; however, while these studies were performed at the genetic level, no studies have established signaling at the protein level^28,30^. This is largely due to the absence of commercially available reagents. In this study, we demonstrate that ActE is a genuine TGFβ ligand that signals through ALK7 with resistance to antagonism akin to its most similar family member, ActC. Modeling suggests that ActE and ActC share similar molecular features (Fig. 1) that define a distinct subgroup within the activin class ligands. Thus, the activin class can be structurally and functionally partitioned into three subclasses ActA/B, GDF8/11 and ActC/E.

Despite current progress, a comprehensive understanding of activin class type I receptor specificity remains incomplete. This is primarily due to activin ligands bearing exceedingly low affinity for the type I receptors which confounds the use of biochemical and biophysical methods to characterize these interactions. Structural information on the different activin ligands shows that the type I receptors contact both chains of the dimeric ligand, specifically at the fingertip and wrist/prehelix regions^10,36^. Both of these regions play roles in determining specificity, as revealed through swapping these domains between ligands to alter receptor selectivity^10,37,45^.

While the structures of ALK4 and ALK5 in complex with a ligand have been resolved, we lack the structure of ALK7 in complex with a TGFβ ligand. With ActE now being added to the group of ligands that can activate ALK7, which includes ActB and ActC, determining common features between the ligands may help to better understand the basis for ALK7-specificity. Interestingly, the fingertip regions between ActB, ActC, and ActE vary minimally (Supp. Table 4) while the wrist/prehelix region is more dramatic (Supp. Table 5). Certainly, actual structural data of ALK7 in complex with a ligand will help to ascertain which ligand residues are important for specificity.

Both ActC and ActE have a divergent sequence at the knuckle region of the ligand. For ActA, ActB, GDF8 and GDF11, the sequence (I/V)AP is highly conserved. In fact, this epitope is further conserved across most of the BMP ligands, including BMPs 2,4,5,6,7 and GDFs 5,6,7. The alanine of the motif points directly into a cluster of hydrophobic resides at the center of the type II receptor interface^7,8,11^. ActC and ActE have a glutamine at the corresponding position.

This difference weakens interaction with the type II receptor indicating that the other activin class molecules would readily out compete ActC and ActE for binding. Interestingly, this modification in ActC also weakens interactions with the extracellular antagonist, which typically bind the other activin class ligands with very high affinity^11^. Thus, one would expect in an environment without antagonists, signaling from activin class ligands that possess high affinity for the type II receptors would dominate over ActC and ActE. However, in the presence of an extracellular antagonist such as FS288, which would block the high affinity type II binding ligands, ActC or ActE signaling may dominate. This might be particularly relevant in the liver which expresses, under certain conditions, the ligands ActA, ActC, and ActE along with follistatin^15,18,46^.

In recent years, there has been an increasing interest in understanding heterodimers within the greater TGFβ superfamily. Unlike the well-studied homodimers, heterodimers of the activin class are poorly understood. Currently, only two activin heterodimers, ActAB and ActAC, have been shown to have biological activity^47–49^. However, several activin heterodimers, ActAE, ActBC, and ActCE, have been detected through overexpression studies *in vitro*^15,17,50,51^.

Compared to their homodimeric counterparts, heterodimers are unique in that they can utilize a larger array of receptors^11,52^. Furthermore, heterodimers can possess different binding capabilities for both receptors and antagonists. For example, ActAC, can signal through both ALK4 and ALK7 and exhibits moderate resistance to follistatin antagonism compared to homodimeric ActA which signals only through ALK4^11^. Unfortunately, due to difficulty in detection, production, and purification of recombinant heterodimeric proteins, teasing out the biological presence/relevance of these molecules *in vivo* remains unclear. The genes encoding ActA, ActC, and ActE (*Inhba, Inhbc, and Inhbe*) are expressed in rodent liver^15^. This expression pattern allows for the possible production of ActAC, ActAE, and ActCE heterodimers. Whether these heterodimers are relevant and play key physiologic or pathophysiologic roles remains unclear; however, given that ActE homodimers can signal, it is now important to consider the role of ActE heterodimers in different biological settings.

Recently, the type I receptor, ALK7, has been heavily implicated in altering fat metabolism in adipose tissue. While ALK4 and ALK5 expression is generally widespread, ALK7 is predominately expressed in adipose tissue but is also detected in the pancreas, brain, and reproductive tissues^44,53–55^. Because ALK7 is primarily expressed in adipose tissue, studies have mainly focused on understanding its role in increasing fat accumulation. ALK7-deficient mice were shown to have increased lipolysis and resistance to high fat diet-induced obesity (DIO)^56–58^. DIO mice treated with the ALK7-specific neutralizing antibody also showed decreased adiposity through enhanced lipolysis^59^. These results, though interesting, do not implicate specific TGFβ ligands in driving these phenotypes, as GDF3, ActB, and ActC all use ALK7 as a signaling receptor.

The data here now indicates that ActE is also an ALK7-specific ligand and should be considered as a possible physiological agonist of ALK7 in adipose tissue. This idea is supported by two recent human-exome sequencing studies designed to identify genes linked to abdominal obesity, along with the recent mouse studies looking at the ligands possible role in fat mobilization^31–33^. Individuals with loss-of-function *INHBE* mutations or variants had favorable fat distribution and metabolic profiles, such as lower triglycerides and higher levels of HDL cholesterol. Interestingly, variants of the ALK7 (*ACVR1C*) gene in humans leads to similar phenotypes (favorable fat distribution and protection from type 2 diabetes) as those shown in variants of the *INHBE* gene^60^.

A recent mouse study by Regeneron showed that *Inhbe* overexpression may down regulate adipocyte lipolysis through the type I receptor ALK7^33^. Blockade of ALK7 using a neutralizing antibody ameliorated the phenotype caused by INHBE overexpression. Since *Inhbe* overexpression could impact other activin signaling molecules, whether these phenotypes are due to increased levels of homodimeric ActE or ActE containing heterodimers is still unknown. It is possible that overexpression of *Inhbe* has a broader impact on the activin family of ligands. While the current study establishes a direct connection of the Act E ligand to the type I receptor ALK7.

Interestingly, overexpression of *Inhbe* led to an increase in the expression of Fstl3, a known activin antagonist^33^. Because we established that ActC and ActE are resistant to antagonism by follistatin family antagonists, this may suggest that the expression of Fstl3 in adipose tissue would block signaling from other activin ligands (ActA, ActB, GDF8, and GDF11) and allow ActC or ActE signaling to dominate. Mechanistically, this may provide a means for ActC and ActE, which have low affinity for the type II receptors ActRIIA and ActRIIB to signal in the presence of ligands which have high affinity for the type II receptors. Thus, Act E signaling might trigger a feed forward mechanism where ActE upregulates Fstl3 to suppress other activin class molecules, while ActE would be resistant to this antagonism. Whether this mechanism plays into the regulation of fat metabolism must be investigated further. Altogether, our results showing that ActE signals through ALK7 provide a connection to these genetic studies and implicates ActE as a hepatokine that may play a direct role in the liver-adipose axis.

Overall, our results show that ActE is a new member of the activin class that can signal like a canonical TGFβ ligand to stimulate the SMAD2/3 pathway. Like ActC, ActE is specific for ALK7 and utilizes the typical type II receptors, ActRIIA and ActRIIB, and has the ability to escape canonical extracellular antagonists. This work will allow further studies to address the biological role of ActE homodimer (or potential heterodimers) signaling through ALK7 in the liver and adipose tissue.

### Experimental Procedures Structural analysis and modeling

AlphaFold 2.3.0 was used to create models of ternary complexes of ActA, ActB, ActC, and ActE with the extracellular domains (ECDs) of the receptors, ActRIIB, ALK4 or ALK7. Five AMBER relaxed models were generated for each complex and the top ranked model was used in further analysis. Input FASTAs for the ligands and receptor ECDs were aligned and truncated to the amino acid coverage of PDB: 7OLY and 7MRZ^36^. For a more consistent comparison, symmetry mates forming the signaling complex were generated for PDB: 7OLY using PyMOL. Potential interactions were analyzed using PDBePISA and visualized using PyMOL. RMSD comparisons were generated using the alignment tool within PyMOL. The “finger” region was defined as the amino acids between the 2^nd^ and 5^th^ and the 7^th^ and 9^th^ cysteine residues of the mature ligand. The “wrist/prehelix” region was defined as the amino acids between the 5^th^ and the 7^th^ cysteine residues of the mature ligand.

### Construct Design

For production of recombinant human activin E in mammalian cells, the full-length coding sequence of the human DNA was used in the vector pRK5. Flag (DYKDDDDK) and myc tags (EQKLISEEDL) were either placed at the N-terminus (Construct I) or the N-terminus of the mature ligand (Construct II). All constructs have a mutated furin cut site in which the cut site has a single point mutation, A233R. The designed sequences were purchased from Synbio Technologies where the gene was subcloned between EcoRI and XbaI restriction sites.

### Protein expression and purification

#### ActA, ActC, and GDF11

Mature recombinant human ActC was purchased from R&D Systems (Cat. No. 1629-AC-010/CF). Recombinant ActA and GDF11 were expressed and purified a previously described^9,10,36^. In short, conditioned media was collected from CHO DUKX cells that stably express ActA. ActA conditioned media was mixed with an affinity resin made with an ActRIIA-construct (Acceleron). The pH of the resin was lowered to pH 3.0 to dissociate the prodomain-ligand complex. Subsequently, the pH was raised to pH 7.5 and incubated for 2 hours at room temperature. For elution, 0.1M glycine pH 3.0, which was then further separated on a Phenyl Sepharose column (Cytiva) and eluted with 50% acetonitrile/water with 0.1% trifluoracetic acid (TFA). Finally, ActA was placed over a reverse phase C4 column (Vydac) and eluted by a gradient of water/0.1% TFA. Conditioned media produced with a construct containing GDF8 prodomain and GDF11 mature (8Pro/11Mature) was provided by Elevian. Conditioned media was loaded over Ni Sepharose excel resin (Cytiva) and eluted with 20mM Tris pH 8.0, 0.5M NaCl, and 500mM imidazole. For further separation, GDF11 was placed over a reverse phase C18 column and eluted with a gradient of water/0.1% TFA. Finally, protein was dialyzed into 10mM HCl and concentrated.

#### ActRIIA-Fc and ActRIIB-Fc

Recombinant human ActRIIA-Fc and recombinant human ActRIIB-Fc were purchased from R&D Systems (Cat. No. 340-R2-100/CF and Cat. No. 339-RB-100/CF, respectfully).

#### Neutralizing antibodies toward ActRIIA, ActRIIB, ActRIIA/ActRIIB and ALK7

All neutralizing antibodies were provided by Acceleron Pharma, Inc. and prepared as previously described^11^. Briefly, anti-ALK7 was expressed transiently in ExpiCHO cells (Thermo Fisher) while the rest were produced in stable CHO cell lines. Anti-ActRIIA was obtained through phage-display technology, while anti-ActRIIB, anti-ActRIIA/ActRIIB, and anti-ALK7 were produced through Adimab’s antibody discovery platform. The conditioned media was purified on Mab SelectSure Protein A (Cytiva) which was followed by ion-exchange chromatography.

#### FS288 and WFIKKN2

FS288 and WFIKKN2 2 were prepared as previously described^37,39,43^. FS288 and WFIKKN2 were expressed from a stably transfected CHO cell line. FS288 conditioned media was placed over a Heparin Sepharose column (Abcam) in 100 mM NaBic pH8.0 and 1.5M NaCl and was eluted with a low salt gradient. Next, FS288 was placed over a cation exchange protocol over a Sepharose fast flow (Cytiva) in 25 mM HEPES pH 6.5, 150 mM NaCl with a high salt elution gradient. Finally, FS288 was purified over an HPLC SXC column in 2.4 mM Tris, 1.5 mM imidazole, 11.6 mM piperazine pH 6.0 and eluted with a high salt pH 10.5 gradient. WFIKKN2 conditioned media was placed over a Butyl Sepharose column and eluted with 20 mM Phosphate buffer pH 7.4 containing 1 mM EDTA. Subsequently, WFIKKN2 was applied to a MonoQ 10/100 GL column in 20 mM Tris pH 8.0, 25 mM NaCl and eluted with linear salt gradient. For final purification, WFIKKN2 was subject to S2000 size-exclusion chromatography in 20 mM HEPES pH 7.4 and 250 mM NaCl and eluted with a salt gradient.

### Activin E Expression and Conditioned Media Collection

ActE constructs were expressed using ExpiCHO-S cells according to the manufacturer’s protocol. Cells were incubated at 37 degrees Celsius with ≥80% relative humidity and 8% CO2 on an orbital shaker platform. Cells were split to a density of 2.0 x 10^6^, 24 hours pre-transfection and were transfected using a concentration of 1 µg/mL of DNA on the day of transfection.

ExpiCHO enhancer and feed were added to the cells 24 hours post transfection. Media was collected three days post-transfection where cells were pelleted, and media was removed and treated EDTA and PMSF (1mM). Conditioned media was concentrated to a final concentration of 20X using centrifugation filter units (Millipore™ Amicon™) for activity assays.

### Activin E Purification

Briefly, media conditioned with ActE flag-tagged at the N-terminus of the mature ligand was incubated with Anti-DYKDDDDK Affinity Resin (ThermoFisher, Cat No. A36801) at 4℃ with gentle rocking. The resin was washed three times with 20mM HEPES, 150 mM NaCl, and 1mM ETDA. Subsequently, ActE was eluted from the resin with 100 mM Glycine pH 2.0, 150 mM NaCl, 0.5% CHAPS with Pierce™ protease inhibitor cocktail, EDTA free (ThermoFisher, Prod # 88266). Finally, fractions were pooled, concentrated, and dialyzed into 10 mM HCl.

### Western Blot

Media samples were subjected to SDS-PAGE (15% SDS gels) under both non-reducing and reducing conditions for detection of both the dimer and monomer components of the ligand. Gels were then incubated in SDS buffer containing BME and DTT reductants (1M) to reduce proteins in gel. This allows for an increase in anti-body binding prior to transfer to nitrocellulose membranes. After transfer, membranes were blocked in TBS-Tween with 5% milk for 1 hour at room temperature. Membranes were probed with anti-DYKDDDDK-HRP (R&D Systems, Cat. No. HAM85291, 1:2000), anti-myc (9E10, Cat. No. CRL-1729, ATCC, RRID:AB_10573245, 1:10), and anti-INHBE (Novus Biologicals, Cat. No. H00083729-B01P, 1:500) over night at 4 degrees. The secondary anti-body used was goat anti-mouse IgG-HRP (Sigma-Aldrich, Cat. No. DC02L, 1:2500). Flag + control used was Flag-BAP fusion protein (Sigma-Aldrich, P7582).

Membranes were imaged using the SuperSignal West Pico detection reagent (ThermoFisher) per manufacturer instructions and detected using a C-DiGit blot scanner (LI-COR).

### Luciferase reporter assays

#### Transient transfection of ligands

Assays using HEK-293T-(CAGA)12 cells were preformed similarly to previously described methods^9,10,36^. Cells were plated in a 96-well format on poly-D-lysine coated plates at a density of 3 x10^4^ cells/well and grown for 24 hours in serum-containing medium. Cells were transfected with 100ng total of DNA containing either empty vector (EV) and ligand (ActA [pRK5], ActE [pRK5], ActC [pcDNA4] and GDF11 [pRK5]), or EV, ligand, and type I receptor constructs, resistant to inhibition by the small molecule SB-431542 (ALK4-ST, ALK5-ST, and ALK7-ST).

Transfection was carried out utilizing Mirus LT-1 transfection reagent according to the manufacturer’s protocol. Following 24 hours transfection incubation, media was replaced with serum-free media with 0.01% BSA (SF^BSA^, ThermoFisher) with or without 10μM SB-431542 and incubated for 18-20 hours. Cells were then lysed, and luciferase activity was measured using a Synergy H1 hybrid plate reader (BioTek - Winooski, VT).

#### Conditioned media activity

As previously mentioned, cells were plated in a 96-well format on poly-D-lysine coated plates at a density of 3 x10^4^ cells/well and grown for 24 hours in serum-containing medium. For all activity assays, growth media was removed and swapped with serum free media with 0.01% BSA (SF^BSA^, ThermoFisher) and the desired ligand/media samples where two-fold serial dilutions were performed for the conditioned media samples. Purified recombinant protein was added at a concentration of 5nM (ActA, GDF11, ActC) and concentrated media was added at a concentration of 20x (1µL/well). After 18 hours of incubation, cells were lysed and luciferase activity was measured using a Synergy H1 hybrid plate reader (BioTek - Winooski, VT). For assays utilizing ALK4-ST, ALK5-ST and ALK7-ST, 10ng of type I receptor DNA and 40ng of empty vector DNA (50ng total/well) was transfected in 24 hours post-plating using Mirus LT-1 transfection reagent. Each construct contains a single point mutation (pRK5 rat ALK5 S278T (ST), pcDNA3 rat ALK4 S282T, pcDNA4B human ALK7 S270T) within the kinase domain that leave the receptor resistant to inhibition by the small molecule SB-431542. Media was then removed and replaced with SF^BSA^ with 10μM SB-431542 and the desired ligand/conditioned media for an 18-20 hour incubation. For assays that utilized neutralizing antibodies (2.5µg/mL for type II and 7.5µg/mL for ALK7), ActRIIA-Fc, ActRIIB-Fc (6.25, 12.5, 25nM), FS288 or WFIKKN2 (12.5, 25nM), the proteins were added to the ligands in test medium and incubated for 10 minutes prior to addition to the cells. The data was imported into GraphPad Prism and analyzed with unpaired t-test or One-Way ANOVA with multiple comparisons on the means of each individual experiment.

### Adipocyte isolation, differentiation, and treatment for western blot

Adipocyte stem cells were isolated, cultured, and differentiated as previously described^11,61^. Briefly, inguinal adipose tissue was harvested aseptically from male mice (3–4 weeks old) and placed in sterile PBS, followed by mincing and collagenase digestion (1 mg/ml) for 1 hr at 37 °C. Then, the digestion was filtered through a 70 µm mesh and centrifuged to separate the SVF. Following aspiration, the SVF was resuspended in DMEM supplemented with 10% FBS and Pen-strep-amphotericin (Wisent Inc cat. No: 450–115-EL, Saint-Jean-Baptiste, Canada) and plated in a 6-well format at ∼320,000 cells/well. Following expansion over 4 days, cells were differentiated over the course of 4 days using a solution of 5 µM dexamethasone, 0.5 mM 3-isobutyl-1-methylxanthine, 10 µg/ml insulin and 5 µM Rosiglitazone. Adipocytes were then maintained for six additional days prior to experimentation in DMEM/FBS + insulin.

Differentiated adipocytes from SVF cells were then starved in serum-free media for 1 hr, after which they were treated with serum-free media containing ActB (2nM), ActC (2 nM), ActE (20x), or Empty Vector (EV) (20x) for 1hr ± Fst-288 (800 ng/ml). In another set of experiments, differentiated-SVF cells were treated with ActC (2nM), ActE (20x), or EV (20x) for 1hr ± anti-ALK7 antibody (7.5 µg/mL). Concentrations were selected based on in vitro cell-based assays. At the end of the treatments, cells were lysed using RIPA buffer containing protease inhibitors and western analysis was performed using anti-pSMAD2 (Cell Signaling, 138D4, Danvers, MA) or anti-SMAD2/3 antibodies (Millipore, 07–408, Burlington, MA).

## Data Availability

All data are contained within the manuscript.

## Acknowledgements and funding sources

We thank Acceleron Pharma for providing the neutralizing antibodies targeting ALK7, ActRIIA, ActRIIB, and ActRIIA/ActRIIB. We also give thanks to members of the Thompson laboratory for helpful feedback and discussion regarding this manuscript. This work was supported by funding from Alnylam Pharmaceuticals, along with NIH R35GM134923-04.

---

**Supplementary Table 1.** PDBePISA analysis of the type II interface (ActRIIB) of activin ternary models.

| Type II (ActRIIB) |  |  |
| --- | --- | --- |
| Solvent-accessible (Å<sup>2</sup>) |  |  |
| Ligands | Interface | Total |
| ActB | 738 | 7641 |
| ActC | 779 | 7635 |
| ActE | 751 | 7408 |
| ActA (PDB: 7OLY) | 680 | 7726 |

**Supplementary Table 2.** PDBePISA analysis of the fingertip region of the type I interface (ALK4 or ALK7) of activin ternary models.

| Type I Fingertips (Chain A) |  |  |
| --- | --- | --- |
| Solvent-accessible (Å<sup>2</sup>) |  |  |
| Ligands | Interface | Total |
| ActB | 401 | 7651 |
| ActC | 447 | 7635 |
| ActE | 375 | 7408 |
| ActA (PDB: 7OLY) | 346 | 7726 |

**Supplementary Table 3.** PDBePISA analysis of the prehelix/wrist region of the type I interface (ALK4 or ALK7) of activin ternary models.

| Type I Prehelix/wrist (Chain B) |  |  |
| --- | --- | --- |
| Solvent-accessible (Å<sup>2</sup>) |  |  |
| Ligands | Interface | Total |
| ActB | 785 | 7641 |
| ActC | 837 | 7599 |
| ActE | 781 | 7386 |
| ActA (PDB: 7OLY) | 617 | 7726 |

**Supplementary Table 4.** Root mean square difference (RMSD) measurements of the fingertip regions of the type I interfaces.

| Fingertip Region (Chain A) |  |  |  |
| --- | --- | --- | --- |
|  | ActB | ActC | ActE |
| ActB | 0 | 0.19 | 0.231 |
| ActC | 0.19 | 0 | 0.212 |
| ActE | 0.231 | 0.212 | 0 |
| ActA (PDB: 7OLY) | 0.364 | 0.401 | 0.396 |

**Supplementary Table 5.** Root mean square difference (RMSD) measurements of the prehelix/wrist regions of the type I interfaces.

| Prehelix/Wrist Region (Chain B) |
| --- |

|  | **ActB** | **ActC** | **ActE** |
| --- | --- | --- | --- |
| **ActB** | 0 | 0.86 | 0.808 |
| **ActC** | 0.86 | 0 | 0.195 |
| **ActE** | 0.808 | 0.195 | 0 |
| **ActA (PDB: 7OLY)** | 1.047 | 0.659 | 0.761 |

